# False-positive neuroimaging: Undisclosed flexibility in testing spatial hypotheses allows presenting anything as a replicated finding

**DOI:** 10.1101/514521

**Authors:** YongWook Hong, Yejong Yoo, Jihoon Han, Tor D. Wager, Choong-Wan Woo

## Abstract

Hypothesis testing in neuroimaging studies relies heavily on treating named anatomical regions (e.g., “the amygdala”) as unitary entities. Though data collection and analyses are conducted at the voxel level, inferences are often based on anatomical regions. The discrepancy between the unit of analysis and the unit of inference leads to ambiguity and flexibility in analyses that can create a false sense of reproducibility. For example, hypothesizing effects on “amygdala activity” does not provide a falsifiable and reproducible definition of precisely which voxels or which patterns of activation should be observed. Rather, it comprises a large number of unspecified sub-hypotheses, leaving room for flexible interpretation of findings, which we refer to as “model degrees of freedom.” From a survey of 135 functional Magnetic Resonance Imaging studies in which researchers claimed replications of previous findings, we found that 42.2% of the studies did not report any quantitative evidence for replication such as activation peaks. Only 14.1% of the papers used exact coordinate-based or *a priori* pattern-based models. Of the studies that reported peak information, 42.9% of the ‘replicated’ findings had peak coordinates more than 15 mm away from the ‘original’ findings, suggesting that different brain locations were activated, even when studies claimed to replicate prior results. To reduce the flexible and qualitative region-level tests in neuroimaging studies, we recommend adopting quantitative spatial models and tests to assess the spatial reproducibility of findings. Techniques reviewed here include permutation tests on peak distance, Bayesian MANOVA, and *a priori* multivariate pattern-based models. These practices will help researchers to establish precise and falsifiable spatial hypotheses, promoting a cumulative science of neuroimaging.

## Introduction

Along with other fields (Baker, 2016; Collaboration, 2015; Hutson, 2018; Ioannidis, 2005), human neuroscience—and functional Magnetic Resonance Imaging (fMRI) in particular—has been facing a replication crisis. A meta-analysis in 2009 estimated the false positive rates in neuroimaging studies to be up to 40% (Wager et al., 2009). Another recent study suggested that more than 50% of neuroimaging findings are likely to be false positives (Szucs and Ioannidis, 2017). To resolve the current replication crisis in neuroimaging, many researchers have discussed the problems related to small sample size, low statistical power, publication bias, data sharing, and p-hacking (Button et al., 2013; Cremers et al., 2017; Munafò et al., 2017; Nord et al., 2017; Pernet and Poline, 2015; Reddan et al., 2017; Szucs and Ioannidis, 2017; Turner et al., 2018). However, there is an additional important issue related to our common practice in neuroimaging studies—the pervasive practice of presenting hypotheses and research findings in terms of gross anatomical region descriptors—that results in substantial flexibility in testing hypotheses. The problem of flexibility in data collection and analysis have been discussed in other contexts (e.g., researcher degrees of freedom; Simmons et al., 2011), and here we extend these discussions to neuroimaging studies, focusing on spatial models.

In neuroimaging studies, gross anatomical region-level descriptors are commonly used to describe hypotheses and compare current findings with previous ones. For example, we can easily find the following statements in neuroimaging studies: “We hypothesize that [task A] would activate [region X], “ or “We replicated a previous study in which [region Y] was associated with the cognitive [function B].” The problem is that gross anatomical regions, such as amygdala or anterior cingulate cortex, do not have exact voxel-level definitions about their locations and usually contain more than 1,000 voxels (Woo et al., 2014b). There could be tens of thousands of possible patterns that constitute “activation of” a single region. Therefore, hypotheses based on gross anatomical regions subsume thousands of possible ways of finding a positive effect. This permits a high degree of flexibility in determining what can count as a positive finding, which we refer to as “model degrees of freedom.” Unfortunately, current standard mapping approaches and major software packages do not provide any analysis methods to explicitly test which locations and patterns of voxel-level activation should be observed. Without voxel-level specifications and tests of topographical information of activation (e.g., locations and patterns), most hypotheses in existing neuroimaging studies cannot be prevented from being qualitative and exploratory (or hypothesis-generating; Ioannidis, 2005), even though they seem quantitative and confirmatory on the surface. Qualitative and exploratory hypotheses render research findings unfalsifiable and resulting in high false positive rates (Simmons et al., 2011).

The larger the hypothesized region, the worse the problem of model flexibility becomes. For example, as previously shown in Woo et al. (2014b), cluster extent-based thresholding often provides large clusters that cover multiple anatomical brain regions (e.g., Fig. 1B of Woo et al. [2014b] showed an example result in which one cluster contained more than 11 distinct anatomical regions). However, the cluster extent thresholding only tells us that there is “at least one non-null voxel somewhere in the cluster,” which may not be falsifiable at all for large clusters, though the hypotheses could be highly reproducible. The key issue here is low spatial specificity of an implicit spatial model—i.e., poor localization ability and resulting lack of confidence in which brain structure(s) are really activated. It is understandable that researchers prefer methods with high spatial sensitivity to ones with high spatial specificity because sensitive methods can provide better-looking results. In addition, the sizes and shapes of brain regions vary across participants, and registration methods are far from perfect. Therefore, defining exact locations across brains is very challenging. In this sense, region- and cluster-level hypotheses can be useful. However, with these hypotheses, one can conclude that two very distinct maps are replications of one another; for example, two maps may have very distinct patterns of activity or have peak activations in the opposite ends of a large brain structure (e.g., the anterior hippocampus bordering on the amygdala vs. the posterior hippocampus centimeters away, bordering on the caudate tail). Therefore, we need methods to quantify whether two studies activate similar locations or produce similar maps.

**Figure 1.**
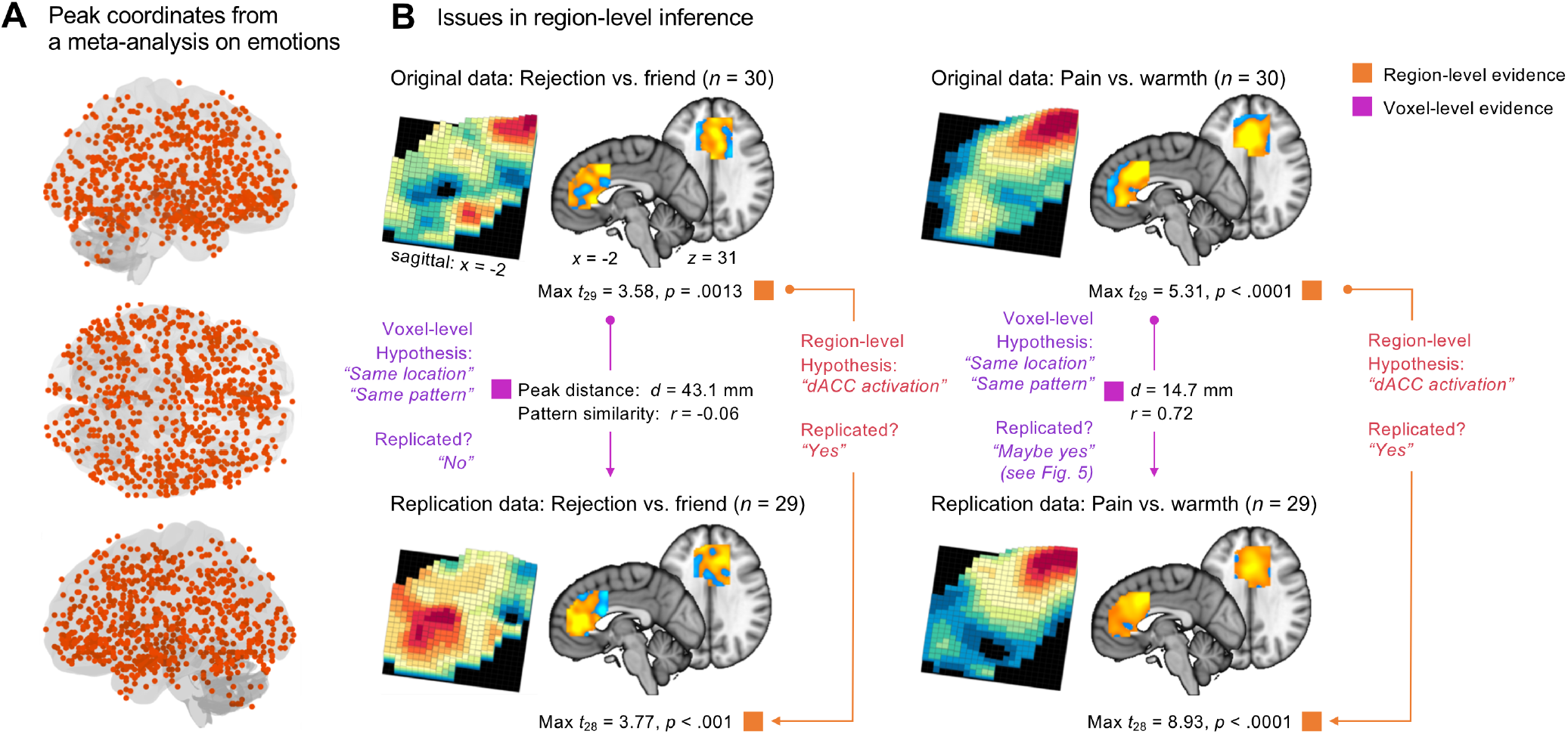
Issues in testing replication using region-level and coordinate-based spatial models. **(A)** Peak coordinates from a meta-analysis for positive and negative emotions (Ashar et al., 2017; Lindquist et al., 2012). The peak coordinates shown here are 856 peaks from 68 fMRI studies that examined aspects of positive or negative emotions, and we only included the studies that reported peak coordinates on the MNI standard space. With these peaks that are distributed all over the whole brain, it is easy to find previous studies that contain peak coordinates near their current findings anywhere in the brain, allowing a post-hoc justification of their findings as replications. **(B)** An illustration of the issues related to using region-level hypotheses in neuroimaging studies. For the illustration, we used an fMRI dataset (*N* = 59; Woo et al., 2014a) that includes somatic pain and social rejection tasks (For more details about the dataset, see **Methods**). To create the original and replication data, we divided the data into two sets while keeping their temporal order of data collection. Then, we used the first dataset as an original study (*n* = 30) and the other set as a replication study (*n* = 29). The figure provides both region-level evidence (orange/red) and voxel-level evidence (violet) for replication and highlights that the region-level and voxel-level evidence can provide opposite conclusions about the replication.

A corollary effect of using region- or cluster-based spatial hypotheses is *low psychological specificity*. One brain region (or even one voxel) typically contains multiple subpopulations of neurons that are functionally distinct (Ito et al., 2003; Kvitsiani et al., 2013; Park et al., 2017) and thus many different tasks and mental processes can activate the same brain region. Therefore, spatial models based on gross anatomical regions cannot achieve a fine-grained understanding of brain-to-function relationships without further specification. For example, the dorsal anterior cingulate cortex (dACC; or anterior midcingulate cortex) is one of the most frequently reported brain regions in the literature (Behrens et al., 2013) with its base rate of significant activation exceeding 20% across tasks and paper topics in human neuroimaging studies (Wager et al., 2016; Yarkoni et al., 2011). It is recruited by many different tasks and mental events including emotional pictures (Ochsner and Gross, 2005), painful stimuli (Wager et al., 2013), emotional pain (Eisenberger et al., 2003), conflict monitoring (Botvinick et al., 1999), prediction error (Hayden et al., 2011), decision making (Kolling et al., 2016), and many others (cf. also see Kragel et al., 2018a; Shackman et al., 2011). Even if these processes activate different subsets of neurons within the dACC and can be distinguished with multivariate patterns of fMRI activity (Kragel et al., 2018a; Krishnan et al., 2016; Woo et al., 2014a), they may all still produce activity in the dACC overall. Therefore, even if an *a priori* hypothesis based on the whole dACC region (e.g., “dACC activation”) is highly reproducible, it is unlikely to provide specific and useful information about brain-to-function mapping. For these reasons, hypotheses based on gross anatomical region descriptors cannot provide a robust foundation for cumulative and reproducible neuroscience; rather, they obscure functional differences and limit their interpretability and falsifiability.

Coordinate-based models, such as a direct comparison of peak coordinates or spherical regions-of-interest around peak coordinates from previous studies, can potentially reduce the level of flexibility in assessing hypotheses and replications by providing tests with better precision compared to region-level models (though they do little to address the issues of functional specificity raised above). However, there are several fundamental limitations in this practice. First, most packages use *ad hoc* algorithms for identifying peak activation locations and provide no inferences about the location or location uncertainty (Kang et al., 2011a; Samartsidis et al., 2017). Most researchers do not specify spatial hypotheses about where brain activations should lie and how uncertain those locations are. This makes testing the ‘replication’ of a hypothesis that was not specified in the original study a somewhat ambiguous venture. In addition, previous studies have shown that peak locations vary widely across different tasks, individuals, and analysis pipelines (Carp, 2012; Kober et al., 2008). For example, peak activations from studies on emotional experience are highly distributed across multiple brain regions (**Fig. 1A**; Kober et al., 2008). Second, coordinate-based models tell us nothing about the patterns of brain activity around the peaks. Two studies with exactly same peak coordinates can have very different patterns of brain activity surrounding those peaks. Third, group-level analyses in fMRI studies usually produce smooth and diffuse brain activation maps (Cremers et al., 2017), which make it intrinsically difficult to locate peak coordinates with certainty and render the coordinate-based models less meaningful. Finally, the use of peak coordinates also provides model flexibility. Currently, there is no consensus on *how close* peak coordinates from two studies should be in order to count as replicates of one another. With the widely spread distribution of peak activations (**Fig. 1A**), researchers can easily find previous studies that contain peak coordinates near their current peak activation *anywhere* in the brain, allowing a spurious, *post-hoc* justification of ‘replicating previous work’. For these reasons, coordinate-based models also cannot provide a solid foundation for the assessment of *a priori* hypothesis and replication.

Multivariate pattern-based models and tests provide a powerful alternative to region- and coordinate-based approaches. Multivariate pattern-based models are increasingly used in fMRI studies to predict behaviors and task parameters due to its high predictive power based on rich information distributed across multiple voxels and regions. Modeling multivariate pattern information is analogous to analysing neural population codes (Kriegeskorte, 2009). A number of studies show convincingly that multivariate pattern-based analysis can capture fine-grained functional information of the brain activity (Alink et al., 2013; Kamitani and Tong, 2005; Shmuel et al., 2010; Swisher et al., 2010) and can more accurately predict perceptions and behaviors than univariate brain maps (Peelen et al., 2006; Woo et al., 2017b; Woo et al., 2014a). More importantly, the pattern-based approach has a potential to provide a precise and quantitative voxel-level specification of the activation locations and the relative levels of activity patterns. For example, a predictive modeling approach (Kragel et al., 2018b; Woo et al., 2017a) aims to develop pattern-based models that are predictive of mental or behavioral outcomes across individuals. These pattern-based models (a.k.a. brain “signatures” [Wager et al., 2013] or “neuromarkers” [Gabrieli et al., 2015]) precisely specify voxel-level weights and define how to integrate new fMRI data from a new individual to produce a single prediction about the outcome. The pattern-based models can serve as *a priori* voxel-level, quantitative, and falsifiable spatial models for testing replications on new brain data from new individuals. This type of multivariate pattern-based models ensures high statistical power because it does not involve any further optimization or multiple comparisons (Gilron et al., 2017; Woo et al., 2017a). It also can remove any wiggle room for further interpretation or redefinition of the models by eliminating any possibility of exploiting ways to find a positive effect.

In the current study, we first illustrate the problems of using gross anatomical region-level descriptors as *a priori* hypotheses with an fMRI dataset from a study comparing somatic pain and social rejection (*N* = 59) (Woo et al., 2014a), highlighting that two very distinct maps of voxel-level tests can be concluded as being replicates of one another based on region-level inferences. Second, from a survey of 135 fMRI studies that claimed a replication of previous findings, we show that a majority of the current fMRI studies (85.3%) rely heavily on region-level hypotheses, and a high proportion of those studies (48.7%) provide no quantitative evidence (e.g., peak coordinates) at all for replication. In addition, when we compared the peak coordinates between the original and ‘replication’ studies, 42.9% of ‘replicated’ findings had peak coordinates more than 15 mm away. Thus, the activation maps that have been counted as ‘replications’ are actually quite different from the original results they claimed to replicate. Third, we highlight the limitation of coordinate-based tests through simulations showing that peak coordinates cannot provide a reliable and stable measure for the underlying patterns of brain activity. The exactly same underlying activation pattern can yield multiple different peaks when noise is added, and two maps that have similar peak coordinates can have distinct activation patterns. Finally, we propose some recommendations for more quantitative testing of hypothesis and replication in neuroimaging studies: (1) Provide quantitative evidence when claiming replications and (2) use explicit and quantitative spatial models and tests, such as permutation tests on peak distance and *a priori* multivariate pattern-based models. These will help to build testable spatial models in neuroimaging and promote the cumulative science of neuroimaging.

## Methods

### Illustration and simulations

To illustrate potential pitfalls of using region-level spatial models, we used an fMRI dataset (*N* = 59) from a previous study (Woo et al., 2014a). The experiment consisted of two tasks: First, in the somatic pain task, participants experienced painful heat (pain condition) or non-painful warmth (warmth condition). In the social rejection task, participants viewed their ex-partner’s photos (rejection condition) or their friends’ photos (friend condition). After we obtained the first-level contrast maps for [pain vs. warmth] and [rejection vs friend] for all subjects, we divided the data into two groups of sequentially acquired participants: An ‘original cohort’ of 30 subjects, and a subsequent ‘replication cohort’ of 29 subjects. We compared their group-level contrast maps for these two cohorts using dACC (anterior midcingulate cortex in particular) as a region-of-interest (ROI). We used the dACC ROI mask from the previous study (Woo et al., 2014a) that showed overlapping activation between the pain and rejection conditions. We additionally smoothed the dACC mask with a 0.5-mm FWHM Gaussian kernel to make the mask large enough for analyzing pattern similarity. We chose to use the ROI test approach to demonstrate the potential pitfalls of the most common approach in the replication studies. We also used the same dataset for a simulation, in which we randomly split the fMRI data into halves (*n* = 30 vs. *n* = 29) 10,000 times and examined whether closely located peak coordinates ensure a high degree of pattern similarity between two group-level maps (**Fig. 4**). Lastly, the same data were used to provide some examples of the recommended methods (**Fig. 5**).

We also conducted another simulation to examine the reliability of peak distance and pattern correlation as a similarity measure of two maps. As shown in **Fig. 3A**, we first created a pair of 100 × 100 matrices (original and replication data) by adding random noise to the underlying ground truth signal pattern (the activity value ranged from 0.2 to 2.0). We combined the signal and the noise to create different levels of signal-to-noise ratio ranging from 0.1 to 1.1. In more detail, the noise was added by adding Gaussian noise with mean = 0 and standard deviation = maximum signal intensity over the ground truth pattern (i.e., 2.0) divided by the desired SNR value. For example, if the SNR was 0.5, the noise was created with random numbers from normal distribution with mean = 0, standard deviation = 4.0; Matlab example, `noise = normrnd(0,2/snr,l00,l00)`. After we smoothed the data, peak distance and pattern correlation were calculated. We repeated this process 10,000 times. **Fig. 3B** shows an example data from one iteration. You can see the simulation code at https://gist.github.com/wanirepo/f34f24e86e49e1f86badfa7c31383e32

### Survey

We surveyed 135 fMRI papers that contain claims of replicating previous findings published between January 2010 and April 2017. To find the papers, we used the following search terms for PubMed database: “fMRI” or “functional magnetic resonance imaging” in the Title/Abstract; “replicate”, “replication”, “replicated”, or “replicates” in the All Fields. We initially acquired 482 papers, which were then filtered with the following exclusion criteria: (1) articles that replicated behavioral results or tasks; (2) articles that used other imaging modalities, such as EEG, PET, or fNIRS; (3) articles that simply mentioned replication (e.g., as a future direction); (4) articles that replicated connectivity studies; (5) review articles; (6) genetic studies. The final number of selected papers was 135.

The 135 replication studies were then categorized into the following seven groups: 1) No specific report: studies that contain no specific spatial information (e.g., peak coordinates or image files to compare) to support their claims of replication. 2) Qualitative region-level comparisons with no report of peak coordinates: studies that used the whole brain search and suggested replication based on qualitative region-level comparisons and did not report peak coordinates. 3) Qualitative region-level comparisons with peak coordinate information. 4) Predefined anatomical ROI test with no report of peak coordinates: studies that used ROIs as prior models and did not report peak coordinates in the paper. 5) Predefined anatomical ROI test with peak coordinate information. 6) Coordinate-based ROI test: studies that used coordinate-based prior models (e.g., 5-mm sphere around a peak coordinate from a previous study). 7) Pattern-based prior models: studies that used multivariate pattern-based prior models.

We also recorded peak coordinates from replication studies (*x_rep_, y_rep_, z_rep_*) if the peak coordinate information was available. We then searched through their original studies that were referenced in the replication studies and extracted peak coordinate information from the original studies if the information was available (*x_orig_, y_orig_, z_orig_*). If the replication cited multiple original studies, we used the peak coordinates that were closest to those in replication studies. Thus, the coordinate distances can be described as assessing the distance to the nearest replicate. Talairach coordinates were converted to Montreal Neurological Institute (MNI) coordinates using the Matlab function, tal2mni.m (Brett et al., 2001). Next, we calculated the Euclidean distance between the peak coordinates using the following equation:

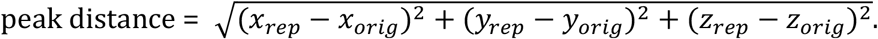

In this manuscript, we use “locations” only in an anatomical sense, as coordinates in standard MNI anatomical space, though in some applications locations can be defined based on brain functional properties as well.

### Recommended analysis methods

In the Discussion, we provide some recommendations to promote the use of quantitative and less flexible spatial models and tests, which include permutation tests for peak distance and pattern similarity, confidence region, multivariate analysis of variance (MANOVA), and multivariate pattern-based classification method.

#### Permutation test for peak distance

A permutation test can be used to compare a peak location in a new study to a fixed reference point from a prior study, i.e., peak distance. For an example analysis, we generated a null distribution of the peak distance between original and replication study data by shuffling the condition labels (in this example, ‘rejection’ and ‘friend’, or ‘pain’ and ‘warmth’) within each participant. The null of the permutation test here posits that no reliable difference exists for the condition contrast (and therefore no reliable peak location) across subjects. The permutation test procedure is as follows: (a) take a fixed peak location from the original study, (b) obtain a peak location from a group-level contrast image (e.g., ‘rejection’ vs. ‘friend’) of the replication study and calculate the peak distance between the original and replication studies, (c) randomly shuffle the condition labels of the replication study (in this case, ‘rejection’ and ‘friend’) and calculate peak distance for each iteration, (d) repeat (c) for multiple iterations (in this example, 10,000 times), and (e) calculate the probability of observing the peak distance between the original and replication studies given the null distribution of permuted peak distance. If the probability of observing the current peak distance is small enough (e.g., *p* < .05), we reject the null hypothesis and conclude that the original and replication studies have peak locations significantly close to each other.

#### Permutation test for pattern similarity

A permutation test can also be used to compare an activation map from a new study to a fixed activation pattern map from a prior study. The null of this permutation test posits that there is no reliable difference between conditions (and therefore no similarity between contrast maps) across subjects. As a measure of the spatial pattern similarity of two maps, we used Pearson’s correlation (*r*):

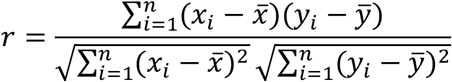

where *n* is the number of voxels included in the map, *x_i_* and *y_i_* are the voxel-level data vectors for two maps (elements are voxels), and 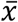 and 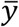 are the mean of the voxel-level data vectors. The permutation test procedure for pattern similarity is similar to the one for peak distance: (a) take a fixed contrast map (e.g., ‘rejection’ vs. ‘friend’) from an original study, (b) obtain a new group-level contrast image from a replication study and calculate the pattern similarity between those two maps, (c) randomly shuffle the condition labels for the replication study (in this case, ‘rejection’ and ‘friend’) and calculate pattern similarity for each iteration, (d) repeat (c) for multiple iterations (e.g., 10,000 times), and (e) calculate the probability of observing the pattern similarity between the original and replication studies given the null distribution of permuted pattern similarity. If the probability of observing the current pattern similarity is small enough (e.g., *p* < .05), we reject the null hypothesis and conclude that the original and replication studies have significantly similar activation patterns.

#### Confidence region

To construct a confidence region based on multiple peak coordinates, we used the method described in Johnson and Wichern (2007; p. 220). The axes of the confidence region in p-dimensional space are defined as:

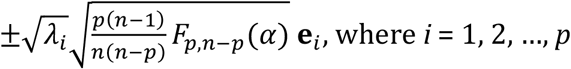

where *n* is the number of peak coordinates, and *p* is the number of dimensions, which is three (i.e., x, y, z) in our case, and *i* is a particular dimension. *F_p,n−p_*(*α*) is the upper (100 *α*)^th^ percentile of the *F_p,n−p_* distribution. In the example case, we construct a 95% confidence region with *p* = 3, and therefore the term should be *F*_3,*n*−3_(0.95). *λ_i_*, and **e**_*i*_, are the eigenvalues and eigenvectors of the sample covariance matrix **S**, respectively, and **S** is defined by

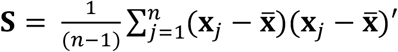

where **x**_1_,**x**_2_,…,**x**_*n*_ are the sample observations, and 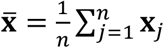.

This has been implemented in the following Matlab functions, conf_region.m and confidence_volume.m (which are available at https://github.com/canlab/CanlabCore and https://github.com/canlab/Canlab_MKDA_MetaAnalysis, respectively). With these functions, one can draw a confidence region using the following lines of Matlab code:

~~~
≫ results = confidence_volume(xyz);
≫ surf(results.xP, results.yP, results.zP);
~~~

#### Bayes Factor calculation for multivariate analysis of variance (MANOVA)

With MANOVA, we can test whether two independent sets of peak coordinates are from the same or different distributions by comparing their multivariate means on the x, y, z space. The null hypothesis of the MANOVA test is that two sets of peak activations are from the same distribution, and it can be rejected when the two sets of peak coordinates are located separately. In the context of testing replication, a Bayes factor provides better tests for confirming replication because it can quantify the likelihood probability of a null hypothesis (i.e., replication success) against an alternative hypothesis (i.e., replication failure) (Rouder et al., 2012; Rouder et al., 2009). We implemented the Bayesian MANOVA and Bayes factor calculation using the BRMS package in R (Bürkner, 2017) and also made a website to provide a web-based Bayes factor calculation at http://cocoanlab.skku.edu/bayes_factor_bayesian_manova. For the accurate calculation of Bayes factors, it is crucial to use the correct priors, and in the implementation, we used weakly-informative priors recommended by Gelman and Hill (2007) Chapters 13 and 17 and STAN manual [v2.17.0] 9.13 and 9.15 (Carpenter et al., 2017). The R-code for the bayes factor calculation is available at https://github.com/cocoanlab/falsepositiveneuroimaging. In this analysis, higher Bayes factors in favor of null hypothesis (BF_01_) provides supporting evidence for replication.

#### Multivariate pattern-based classification

If an *a priori* pattern-based model is available, one can calculate the pattern expression values using dot-product but other similarity metric (e.g. Pearson’s correlation, Spearman correlation, cosine similarity, etc.) can also be used:

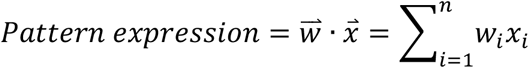

where *n* is the number of voxels within the pattern-based model, *w* is the voxel-level predictive weights, and *x* is the test data. An *a priori* pattern-based model is composed of predictive weights 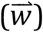 across voxels, specifying locations and patterns of activation. The weights tell us how to integrate fMRI data into a single prediction, which then can be used for classification tests or regression analyses. In our example analysis, we calculated the dot product between the *a priori* pattern-based model for [rejection vs. friend] and [pain vs. warmth] trained on the original data, *n* = 30, and the test image data from the replication data, *n* = 29. Then, we conducted a classification test on the pattern expression values with the forced-choice test and the binomial test to determine whether the observed accuracy is significant.

## Results

### Illustration of the issues related to region-level tests

As illustrated in **Fig. 1B**, when relying only on a region-level hypothesis such as “significant activation within dACC, “ one can easily conclude that the previous findings are successfully replicated even with two very distinct activation maps at the voxel-level. For the [rejection vs. friend] contrast, both original (*n* = 30) and replication (*n* = 29) data contain significantly activated dACC voxels while their peaks are located far away from each other, *d* = 43.1 mm, and the patterns of activations between two maps are uncorrelated, *r* = −0.06. For the [pain vs. warmth] contrast, two maps from the original and replication data show peak activations that are relatively close to each other, *d* = 14.7 mm, and a high degree of pattern similarity, *r* = 0.72 (quantitative tests on these values will be proposed in the **Discussion**). If we make conclusions based only on the region-level evidence, we should conclude that findings for both contrasts, i.e., [rejection vs. friend] and [pain vs. warmth], are successfully replicated, but voxel-level evidence provides a different conclusion for the [rejection vs. friend] contrast, which is not highly replicable at the voxel-level.

The same issue can occur when comparing two different psychological states based on fMRI activation maps. If a researcher relies only on a region-level test, one can conclude that social rejection and physical pain share neural representations within the dACC based on significant activations within the region across two datasets. However, the voxel-level examination comparing the two contrast maps (i.e., one for [rejection vs. friend] and the other for [pain vs. warmth]) suggests a different conclusion. In the first dataset (*n* = 30), two maps had peaks close to each other, *d* = 8.5 mm, and similar activation patterns, *r* = 0.49, supporting shared brain representation across rejection and pain within the dACC. However, in the replication dataset (*n* = 29), the peaks from two maps were located far from each other *d* = 46.0 mm and the activation patterns were negatively correlated, *r* = −0.19, suggesting that pain and rejection do not share neural representations.

### Survey results

As shown in **Fig. 2A**, we found that a majority of the current fMRI replication studies rely heavily on region-level assessment (qualitative region-level comparison and predefined anatomical ROI test): 24.5% and 60.8% of the 135 surveyed studies respectively used the qualitative region-level comparison and the predefined anatomical ROI test for the replication assessment. Note that the currently most popular method (60.8%) for the replication assessment is the predefined anatomical ROI test, which might look like a positive sign for good analysis practice. However, **Fig. 2B** suggests that it is not the case: The qualitative region-level comparison and the predefined anatomical ROI test did not differ in their peak distances between original and replication studies (*t*_44.8_ = 0.57, *p* = .569, two sample *t*-test), suggesting that both approaches are similarly liberal in assessing replication (see below for more detailed comparisons).

**Figure 2.**
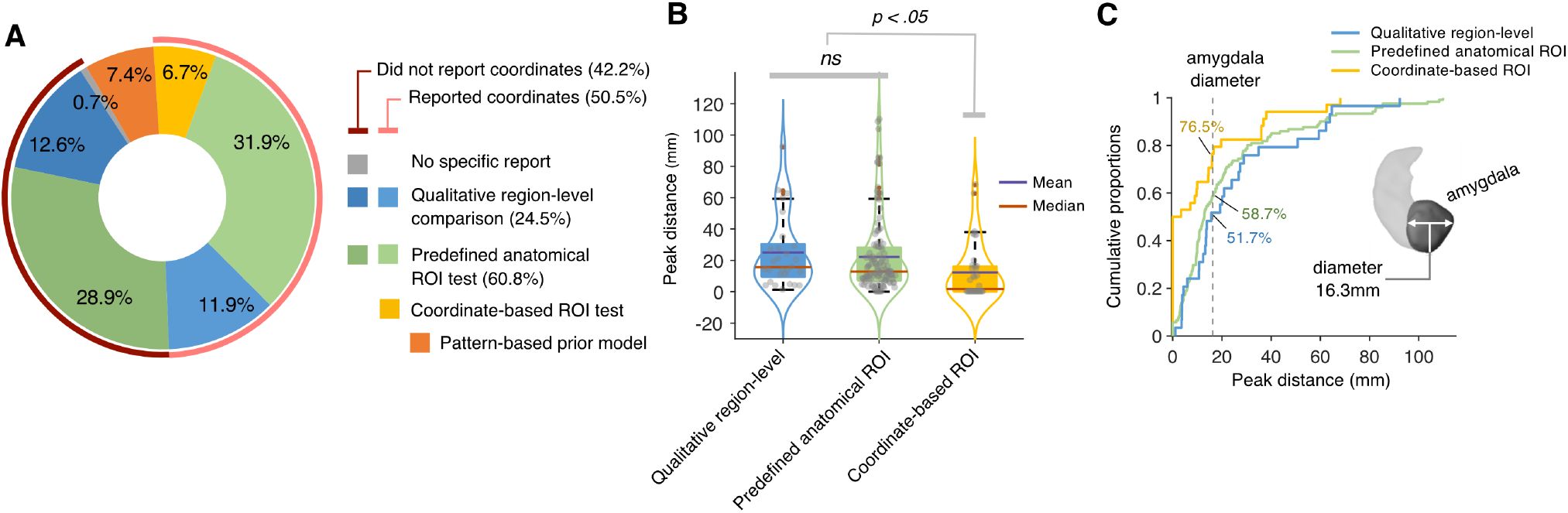
Survey results. We surveyed 135 fMRI papers that contain claims of replicating previous findings and were published between January 2010 and April 2017. **(A)** The pie chart shows the proportions of seven categories for the replication studies. The seven categories are as follows: (1) ‘No specific report’ refers to the studies that include no specific information to support their claims of replication. (2) ‘Qualitative region-level comparison with no report of peak coordinates’ refers to the studies that suggested replication based on qualitative region-level comparisons and did not report peak coordinates. (3) ‘Qualitative region-level comparisons with peak coordinate information’. (4) ‘Predefined anatomical ROI test with no report of peak coordinates’ refers to the studies that used anatomical ROIs as prior models and did not report peak coordinates in the paper. (5) ‘Predefined anatomical ROI test with peak coordinate information’. (6) ‘Coordinate-based ROI test’ refers to the studies that used coordinate-based prior models (e.g., 5-mm sphere around a peak coordinate from a previous study).(7) ‘Pattern-based prior model’ refers to the studies that used *a priori* multivariate pattern-based models. The category groups except for ‘Pattern-based prior model’ were subdivided into two groups based on whether the study reported peak coordinates or not. **(B)** The box-violin plots present the distributions of peak distance (in mm) between replication and original studies for the three study categories that reported peak coordinates. Dark purple and red lines respectively indicate the mean and median values. Studies that used coordinate-based prior models showed significantly shorter peak distance compared to others (*p* < .05, two-sample *t*-test). **(C)** The cumulative distribution plot shows the cumulative proportions of the peak distance values. To provide a benchmark, we used the average diameter of amygdala (16.3mm from Brabec et al., 2010). More than 40% of the studies within the ‘qualitative region-level comparison’ (41.3%) and ‘predefined anatomical ROI test’ (48.3%) categories showed peak distances longer than the amygdala diameter.

Importantly, 42.2% of the 135 replication studies (dark red line outside of the pie chart) did not even report peak coordinates, indicating that these studies claimed replication without any quantitative voxel-level evidence. 6.7% of the studies used a coordinate-based ROI test, in which spatial hypotheses are formed using peak coordinates from previous studies, and 7.4% of the studies used *a priori* pattern-based models.

**Fig. 2B** displays distances between peak coordinates from original vs. replication studies (peak distance) across different study categories. The studies that used the qualitative region-level comparison and the predefined anatomical ROI test showed similarly long peak distances (for the qualitative region-level comparison studies, mean = 25.0 mm, median = 15.7 mm, SD = 23.4 mm; for the predefined anatomical ROI studies, mean = 22.2 mm, median = 12.8 mm, SD = 24.4 mm). By contrast, the coordinate-based ROI test studies fared somewhat better, showing peak distances significantly closer than the other two study categories (mean = 12.2 mm, median = 1.7 mm, SD = 18.1 mm, *t*_62.7_ = −2.87, *p* = .006, two sample *t*-test).

We additionally compared distributions of the peak distances across three groups using the Kullback-Leibler (KL) divergence; lower KL divergence indicates more similar distributions. With the probability density function using the bin size of 4 mm, KL divergence for the qualitative region-level comparison vs. predefined anatomical ROI groups showed smaller KL divergence, KL = 3.84, than the KL divergence for the predefined anatomical ROI vs. coordinate-based ROI groups (KL = 5.47) and for the qualitative region-level vs. coordinate-based ROI groups (KL = 9.13), suggesting that the difference between the qualitative region-level comparison and the predefined anatomical ROI groups was smaller than their differences with the coordinate-based ROI group. The mean and median peak distance across all three groups were respectively 20.8 mm and 12.8 mm with standard deviation of 23.5 mm.

In **Fig. 2C**, we used the amygdala, the average diameter of which is 16.3 mm (Brabec et al., 2010), to provide a reference point for the peak distances of the surveyed studies. 48.3% of the qualitative region-level comparison studies, 41.3% of the predefined anatomical ROI test studies, and 23.5% of the coordinate-based ROI test studies (overall 39.1%) had peak distances longer than the amygdala’s diameter, highlighting the fact that around 40% of the replication studies in the neuroimaging field claimed replication even with peak differences larger than the size of amygdala.

### Limitation of coordinate-based models in evaluating replications

Though assessing peak coordinates between the original vs. replication studies (e.g., peak distance) could serve as a quantitative method for evaluating replications, peak coordinates provide a suboptimal metric, in part because peak locations are not a measure of central tendency and thus suffer from a poor basis of support in the data. The simulation results shown in **Fig. 3C** and **Fig. 4** clearly show the limitations of using peak distance as an evaluation method for replication.

**Figure 3.**
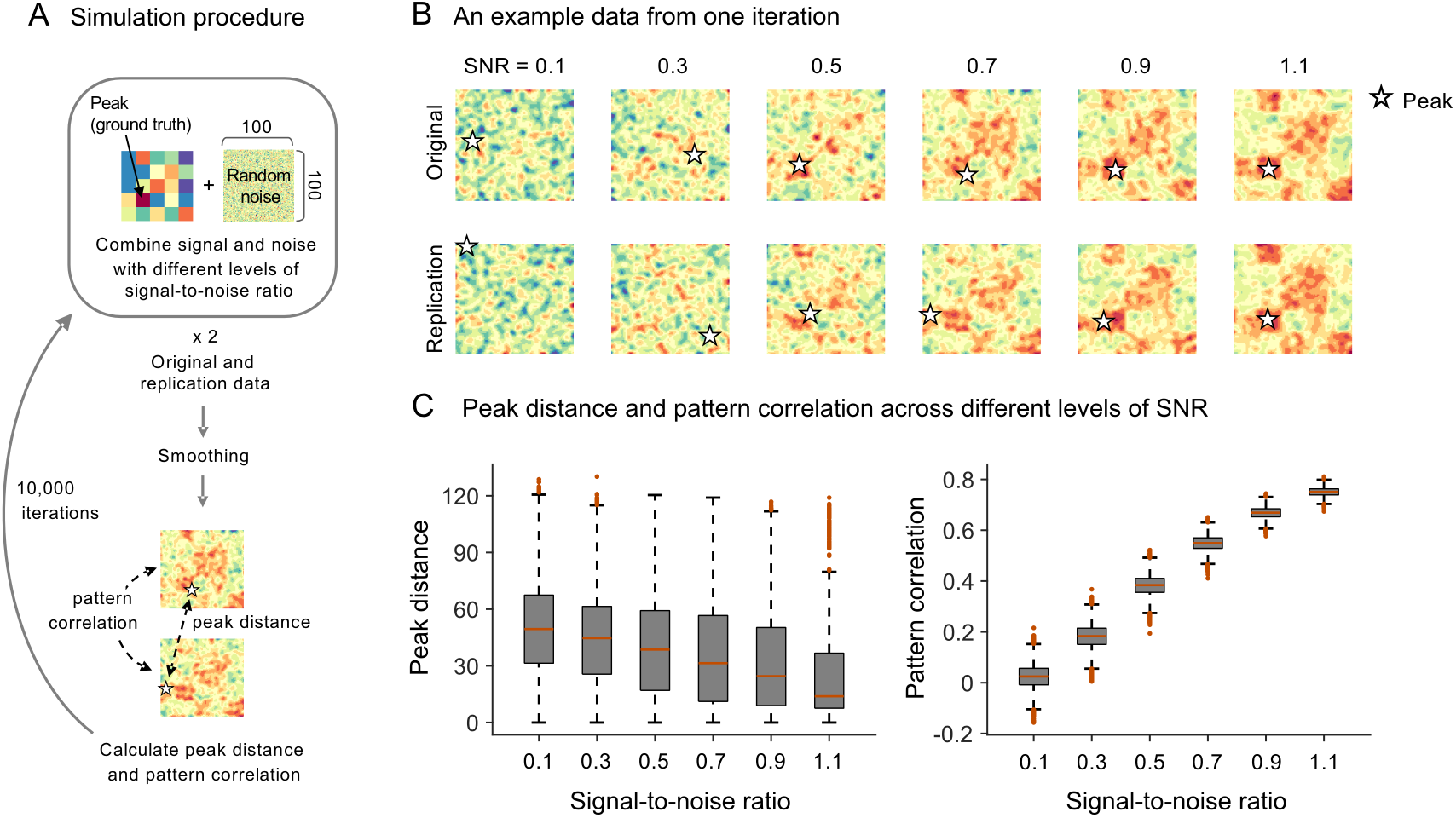
Simulation 1. **(A)** To examine the reliability of peak distance and pattern correlation when comparing two simulated data with the same ground truth patterns of signal, we created a pair of 100 x 100 matrices (original and replication data) by adding random noise to the underlying ground truth signal pattern. The ground truth pattern consists of 5 x 5 patches, each of which is a 20 x 20 matrix with the same activation values ranging from 0.2 to 2.0. We combined the signal and the noise with different levels of signal-to-noise ratio ranging from 0.1 to 1.1. After smoothing, peak distance and pattern correlation between two data matrices were calculated. We did not include two edge rows and columns when detecting peaks to reduce the edge effects due to smoothing. We repeated this process 10,000 times. **(B)** An example data from one iteration. Stars indicate peak locations of the data matrices. **(C)** Distributions of peak distance and pattern correlation across different levels of SNR.

**Figure 4.**
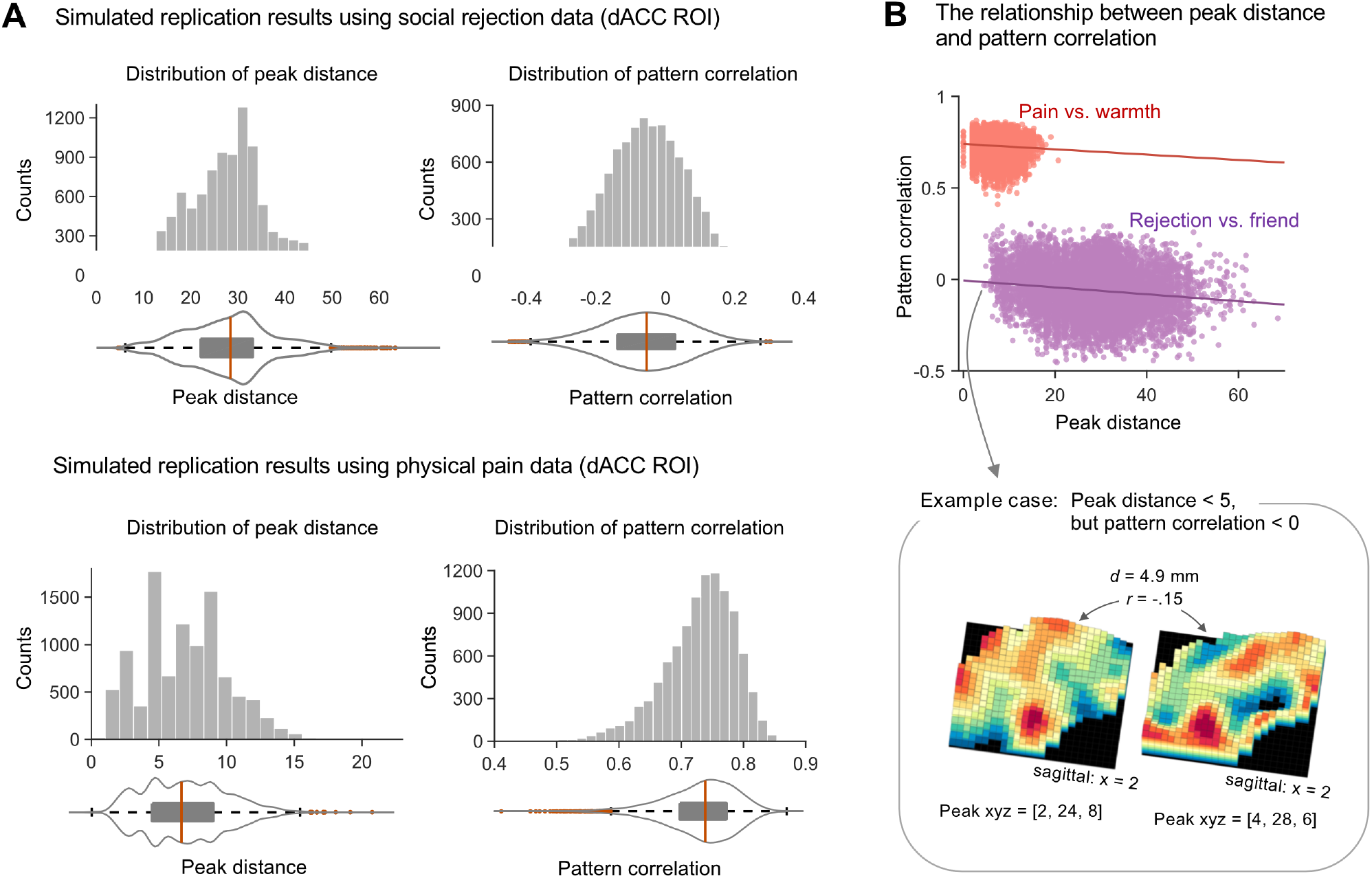
Simulation 2. **(A)** The histograms show the distributions of peak distance and pattern correlation within the dACC between group contrast maps from two randomly half-split datasets using the same fMRI dataset that we used for illustration (Woo et al., 2014a). We treated the first and second sets of the half-split data as original and replication studies, respectively. The top panel shows the results with the social rejection data, [rejection vs. friend], and the bottom panel shows the results with the physical pain data, [pain vs. warmth]. **(B)** The scatterplot shows the relationship between pattern correlation (y-axis) and peak distance (x-axis) for both datasets, red: pain vs. warmth, purple: rejection vs. friend. The reference lines are the least-square lines. The bottom panel displays an example case where the peak distance is very short (*d* < 5 mm), but the patterns of brain activity are distinct (*r* < 0), highlighting the fact that close peaks do not necessarily imply similar patterns of brain activity.

First, peak coordinates cannot provide a reliable and stable measure for the underlying patterns of brain activity because peaks are more vulnerable to noise than multivariate pattern information. The exactly same underlying ground truth activation pattern can yield multiple different peaks when noise is added. For the simulation shown in **Fig. 3**, we created a pair of matrices by combining one ground-truth activation pattern with different random noises at each iteration and compared two data matrices using peak distance and pattern correlation after smoothing. We combined the signal and the noise with different levels of signal-to-noise ratio (SNR) that ranged from 0.1 to 1.1, increasing by 0.2. As expected, the average peak distance was decreased and the pattern correlation was increased as the SNR increased. However, the peak distance values were highly variable across all levels of SNR, whereas the pattern correlation values had much lower variance than peak distance. To quantify this, we compared the effect sizes of SNR increases on peak distance and pattern correlation using Cohen’s *d* (a mean difference between two adjacent levels of SNR divided by pooled standard deviation). Peak distance showed small effect sizes; absolute Cohen’s *d* for the decreases of peak distance ranged from 0.19 to 0.23 with mean *d* = 0.21. In contrast, pattern correlation showed large effect sizes ranging from 1.71 to 1.83 with mean *d* = 1.80.

Second, we did a simulation with real data and found that closely located peaks cannot ensure the high degree of pattern similarity. As shown in **Fig. 4**, we compared peak distance and pattern similarity within the dACC between two group-level contrast maps for [rejection vs. friend] constructed from 10,000 iterations of random split-half samples. If the maps are reproducible, two contrast maps from split-half samples should show closely located peaks (i.e., short peak distance) and high degree of pattern similarity. The results showed that the maps for the contrast of [rejection vs. friend] are not highly reproducible: The average peak distance over 10,000 iterations was 27.9 mm and the mean pattern similarity was *r* = −0.058. A linear regression analysis with peak distance as a predictor and pattern correlation as an outcome showed that peak distance was a weak, but significant, negative predictor for pattern correlation 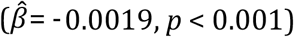. However, the peak distance explained only 2.03% variance in the pattern similarity values. In addition, the model intercept was negative (intercept = −0.006) suggesting that a peak distance close to 0 cannot guarantee a positive pattern correlation. **Fig. 4B** highlights an example case where the peak distance between two half-split data was very short (*d* = 4.9 mm), but still showed negative pattern correlation (*r* = −0.15). Therefore, it is possible that two brain maps with very different brain activation topography can be considered to be same if we examine only peak coordinates.

We also conducted the same simulation for the [pain vs. warmth] contrast. As shown in the bottom panel of **Fig. 4A**, the [pain vs. warmth] contrast showed a significantly lower average peak distance (6.9 mm) compared to the [rejection vs. friend] contrast (27.9 mm). This suggests that peak distance could be an adequate measure in some cases (e.g., when the spatial variability is low). However, the linear regression results using the simulated data for [pain vs. warmth] were quite similar to the one for [rejection vs. friend]: Peak distance was a weak predictor of pattern similarity, 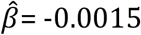 and was able to explain only 0.6% variance in pattern similarity, supporting the idea that peak distance is not a good predictor of pattern similarity.

## Discussion

### Region-level models allow flexibility in determining what can count as positive findings

From the survey of 135 fMRI studies that contain claims about replication of previous studies, we found that the currently most popular spatial models are region-level models (85.3%; **Fig. 2A**). This is important to note because, with these models, many combinations (likely thousands) of different activation patterns can be interpreted as positive findings of a region-level hypothesis, such as “amygdala activity.” Anatomical brain regions usually contain more than 1,000 voxels, and thus the actual number of hypotheses becomes the number of possible combinations among voxels within the regions. In other words, when we test a hypothesis based on a gross anatomical region, we are testing not just the “cover” hypothesis but also a large number of unspecified “hidden” hypotheses, that can all lead to positive findings. The unspecified sub-hypotheses make the cover hypothesis less falsifiable, leading to false positive findings and replication failure in the long run. In addition, it is difficult to establish sensitive and specific links between brain measures and mental categories—a central goal of cognitive neuroscience—without more precise specification of which voxels and patterns (i.e., relative values across voxels) should be activated.

This concern is well supported by our survey results. The studies that employed qualitative region-level comparisons or predefined anatomical ROI tests showed a peak distance to the nearest claimed “replicating finding” greater than 22.6 mm on average, and around 40% of these studies had peak distances greater than the amygdala’s average diameter (16.3mm), suggesting that region-level tests indeed allowed presenting very different activation maps as being replicated. This was only among studies reporting peak distances; almost half (42.2%) of the surveyed studies did not even report quantitative voxel-level (peak or pattern) evidence for replications.

To reduce false positives and the false sense of reproducibility in neuroimaging studies, researchers need to develop and use formal statistical analysis methods to support quantitative and explicit spatial models and hypotheses. These methods should provide statistical inferences about where brain activations are located and how uncertain those locations are. Then, tests of replication should be based on these quantitative spatial models and hypotheses. As a step towards developing quantitative assessment of replications and spatial hypotheses, here we propose some specific recommendations: 1) Provide quantitative voxel-level evidence (i.e., peak or pattern) when claiming replications and 2) use explicit spatial models and tests. If peak locations are the only information available, we recommend using explicit models and tests for peak coordinates, such as permutation tests on peak distance and Bayesian MANOVA tests on peak distributions. These could provide statistical inferences about where a peak coordinate is likely to lie and whether two or more conditions activate same or different peak locations. However, as we highlighted through simulations shown in **Figs. 3–4**, peak locations provide only a suboptimal way to evaluate replications, and therefore spatial pattern-based analyses should be a major direction in future replication studies. We explain each recommendation in more detail below, summarize our recommendations in **Fig. 5** and the **Appendix**, and describe several useful test procedures in the **Methods**.

**Figure 5.**
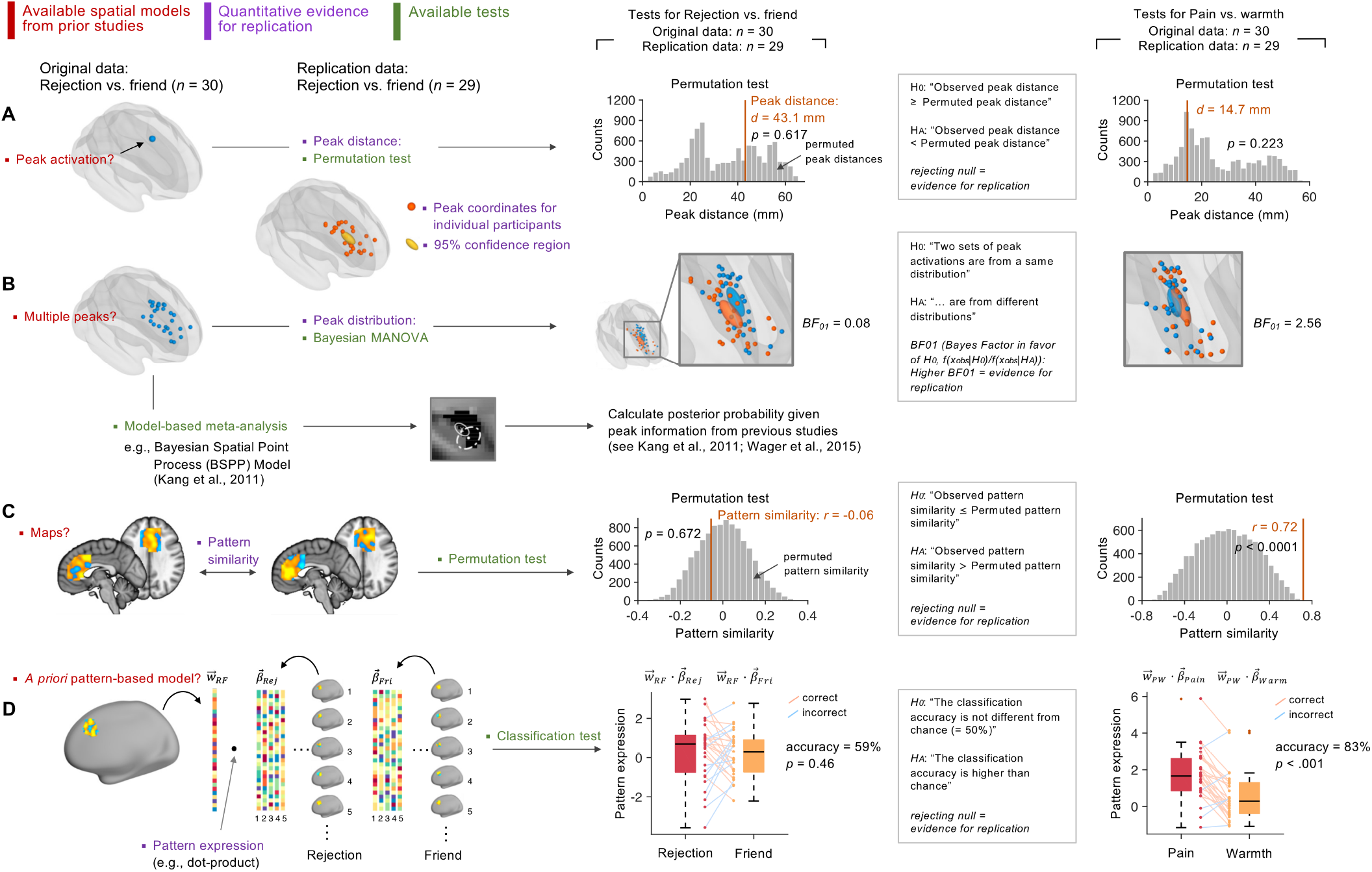
Recommendations. This figure shows our recommendations to reduce flexible and qualitative spatial tests in neuroimaging studies. We provide different options of quantitative voxel-level evidence (violet) and test methods (green) for different types of spatial models available from previous studies (red), ranging from **(A)** when only a small number of peak coordinates are available as spatial models to **(B)** when multiple peak coordinates are available, **(C)** when unthresholded maps are available, and **(D)** when *a priori* pattern-based models are available. For more detailed explanation about each method, please refer to **Methods** and **Discussion**. RF = rejection vs. friend. PW = pain vs. warmth. Rej = rejection. Fri = friend. *H_0_* = null hypothesis. *H_A_* = alternative hypothesis.

### Provide quantitative voxel-level evidence when claiming replications

As proposed in **Fig. 5** (violet font color), one can provide voxel-level evidence for replication using peak coordinates, confidence regions, peak distance, pattern similarity, or pattern expression values. More details about the calculation of these values are included in the **Methods**. To determine which types of voxel-level information to provide, one needs to consider which type of spatial models is available from the original study first. **Fig. 5** provides recommendations and examples for different cases.

When one or only a small number of peak coordinates are available from a previous study, one can provide peak distance along with individual peak coordinates and a 95% confidence region. As our survey results showed, the studies that used the coordinate-based ROI tests reported significantly shorter peak distances than other study categories (**Fig. 2B**), indicating that the coordinate-based ROI tests provide more precise models compared to other region-level tests maybe because the coordinate-based tests have a more limited search space than others. However, going beyond the coordinate-based ROI tests, researchers should be able to report and systematically compare peak coordinates from their current study against the peaks from previous studies using peak distance. If multiple peak coordinates are available from previous studies (e.g., from meta-analyses or from first-level contrast maps), one can provide a peak distribution along with 95% confidence regions for the previous and current studies. A confidence region from the original study could potentially serve as a replication target, and overlapping confidence regions can be an evidence for successful replication. However, one should interpret results with large confidence regions with caution because large confidence regions indicate that the results are spatially variable and have poor spatial specificity. Thus, in addition to whether a study spatially replicates an earlier finding, the size of the spatial confidence region and resulting implications for the precision of localization should be an important consideration.

If unthresholded activation maps or *a priori* pattern-based models are available from previous studies (e.g., from NeuroVault.org [Gorgolewski et al., 2015]), one can use similarity metric, such as pattern similarity and pattern expression values. This provides a quantitative way to measure the similarity of activation patterns across voxels between an original study and a replication study. Such measures are becoming increasingly popular also as measures of representational similarity (Haxby et al., 2014; Kriegeskorte and Kievit, 2013).

### Use quantitative spatial models and tests for peak locations

Though peak locations have many limitations in assessing replication (**Figs. 3–4**) researchers might have no choice other than using peak information. In that case, we recommend using a permutation test or Bayesian MANOVA.

First, when peak distance is used as a voxel-level evidence, one can use a permutation test to examine whether the observed peak distance is shorter than the null distribution of permuted peak distances under null hypothesis. Here, the null hypothesis is that there is no reliable peak location exists for the condition contrast across subjects. The permutation test can resolve the issue of having no standard criterion for how short the peak distance should be to count as replication. For the example analysis (**Fig. 5A**), we generated null distributions of peak distance by shuffling the condition labels (e.g., ‘rejection’ and ‘friend’) within each participant (for more details, see **Methods**). The permuted data showed the intrinsic distribution of the peak distance within the dACC ROI, which was not a normal distribution. The test results showed that the observed peak distance for [rejection vs. friend] (*d* = 43.1 mm, see **Fig. 1B**) was not significantly shorter than the permuted peak distance, *p* = 0.617, suggesting that the replication study failed to reproduce the brain activation map more closely compared to random (null) maps. The test result for [pain vs. warmth] (*d* = 14.7 mm) also suggested replication failure, *p* = 0.223. This may be a function of the poor measurement properties of peak distance, as discussed above, rather than a failure to find reproducible activation, considering other test results presented below.

Second, when multiple peak coordinates are available from previous studies, one can run Bayesian MANOVA or meta-analysis. Bayesian MANOVA can provide evidence for and against the hypothesis that a new study produces peak activation locations consistent with prior studies (a web-based Bayes factor calculation is available at http://cocoanlab.skku.edu/bayes_factor_bayesian_manova). Multivariate confidence regions are useful to visualize the peak distributions around the peak center along with the results of Bayesian MANOVA (for more details of how to construct confidence region, see **Methods**). In our example, as shown in **Fig. 5B**, the Bayes factor in favor of null hypothesis (BF_01_) suggested that the peak distributions of the original and replication studies are distinct for the [rejection vs. friend] contrast, BF_01_ = 0.08. The Bayes factor for the [pain vs warmth] contrast supported null hypothesis, BF01 = 2.56, suggesting successful replication. Note that, similar to the use of confidence regions, one should be cautious about interpreting MANOVA results when peak coordinates are highly variable because high variance indicates that the peak locations are not reliable.

In addition to MANOVA, conducting coordinate-based meta-analysis is strongly recommended to combine multiple peak coordinates (Samartsidis et al., 2017). Meta-analysis provides a principled way of constructing *a priori* hypotheses, preventing an arbitrary cherrypicking of prior studies to favor current findings. Particularly, with model-based meta-analysis such as Bayesian Spatial Point Process (BSPP) model (Kang et al., 2011b; Wager et al., 2015), one can calculate the posterior probability of observing the current data based on the peak information from previous studies. We do not provide a full description of this method in this paper because it is also beyond the scope of the current study.

### Multivariate pattern-based tests

A multivariate pattern-based approach provides a powerful alternative to the region- and coordinate-based approaches in reducing flexibility in hypothesis testing. In addition to its ability to capture information distributed across multiple voxels and regions, the pattern-based approach can provide a precisely defined *a priori* model, which contains specific information about the activation location and relative patterns of activity levels. The pattern-based *a priori* model removes any wiggle room for further interpretation or redefinition of the model, eliminating the possibility of hiding a large number of hidden hypotheses. Researchers can even save their pattern model as an image file (e.g., a nifti file), allowing easy sharing of the model across researchers and laboratories. In addition, the models can be easily applied and tested across studies and datasets with no further modification, minimizing flexibility in testing and replicating effects in new individuals and studies and making the tests confirmatory, falsifiable, cumulative, and transparent.

As shown in **Fig. 5C**, when unthresholded maps are available from the previous studies, one can use permutation tests for pattern similarity between two brain activation maps (defined by Pearson’s correlation coefficients across voxels). In our example analysis, we generated null distributions of pattern similarity between the dACC activation patterns of two group-level contrast maps by shuffling the condition labels within each participant (for more details, see **Methods**). The test results showed that the observed pattern similarity for the [rejection vs. friend] contrast (*r* = −0.06, see **Fig. 1B**) was not higher than the permuted pattern similarity, *p* = 0.672, suggesting that the replication study failed to reproduce the brain activation map. The test result for the [pain vs. warmth] contrast (*r* = 0.72) showed that the observed pattern similarity was significantly higher than the permuted pattern similarity, *p* < 0.0001, suggesting a successful replication.

Finally, **Fig. 5D** provides an example analysis for multivariate pattern-based marker approach. In the example analysis, we trained pattern classifiers (using linear support vector machines) based on the original data (*n* = 30), one for [rejection vs. friend] and the other for [pain vs. warmth]. Then we tested the exactly same classifier models (without any modifications) on the replication data (*n* = 29) (for more detailed test procedure, see **Methods**). The classification accuracy for the [rejection vs. friend] contrast was 59%, which was not different from chance, *p* = 0.46, suggesting a replication failure. Conversely, the classification results for the [pain vs. warmth] contrast showed a significant accuracy, 83%, *p* < 0.0001, suggesting a successful replication.

Despite its advantages, the multivariate pattern-based approach that uses *a priori* pattern-based models also has a limitation: *A priori* pattern models can utilize only the voxel-level information that is consistent and conserved across people and studies. The amount of information conserved across individuals and studies could be small depending on the target mental events, study populations, and even differences in preprocessing pipelines. To address this limitation, researchers can build and test their hypotheses on the representational space, not on the brain space, using methods such as hyper-alignment (Haxby et al., 2011) or representational similarity analysis (Kriegeskorte and Kievit, 2013). However, even for the representational space-based approach, the same caution should be given to make hypothesis testing and replication assessment more falsifiable and transparent.

## Conclusion

Though anatomical region-level descriptors provide the most popular spatial models for testing and replicating previous findings in neuroimaging studies, the gross anatomical region descriptors can be interpreted in many different ways without further specification of the hypothesized locations and patterns of activation. These region-level models introduce unwanted flexibility into a study, resulting in unfalsifiable hypotheses, false positive findings, and replication failure. To build a more cumulative and falsifiable science of neuroimaging, we recommend using more quantitative spatial models in testing and replicating previous findings and reporting them. First, we recommend providing quantitative spatial evidence for the claimed replication. From our survey on 135 studies that suggested replications of previous findings, we found that a high proportion of the studies (42.2%) provided no quantitative evidence for replication at all. Second, if researchers form their *a priori* hypotheses using peak coordinates from previous studies, we recommend conducting formal statistical tests on peak distance or peak distribution using permutation tests, Bayesian MANOVA, or meta-analysis. Lastly, we strongly recommend using the *a priori* multivariate pattern-based approach, which can eliminate flexibility in interpreting *a priori* hypotheses by providing precise definitions of activation locations and relative patterns of activity. These practices will provide researchers with more robust spatial tests, helping us move one step towards resolving the current replication crisis in neuroimaging studies.

## Appendix. Summary of recommendations

## Recommendation 1: Provide quantitative evidence when claiming replications

- Report **peak distance** between the original and replication studies:

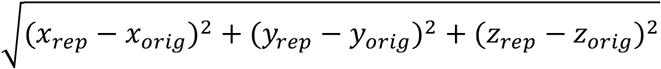
- Construct and visualize **confidence regions** around estimated peak locations:

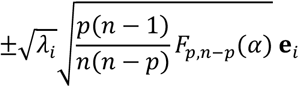
- Estimate **pattern similarity** between the original and replication maps:

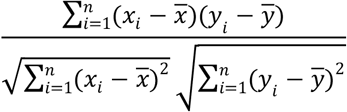
- Calculate and test **pattern expression values** using *a priori* pattern-based models:

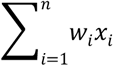

## Recommendation 2: Use spatial models and tests for peak locations

- **Permutation tests for peak distance** Permutation tests can be used to examine whether the observed peak distance between original and replication studies is significantly shorter than the permuted peak distance.
- **Bayesian MANOVA** Bayesian MANOVA can be used to test whether the multivariate sample means of two sets of peak coordinates (e.g., a set of individual participants’ peak coordinates for an original study and a set for an attempted replication) are the same or different using Bayes Factors to quantify evidence for replication.
- **Coordinate- or model-based meta-analysis** These methods can be used to estimate the posterior probability of observing the current data based on the peak information from previous studies.

## Recommendation 3: Use *a priori* multivariate pattern-based models

- **Permutation tests for pattern similarity** Permutation tests can be used to examine whether pattern similarity of activation maps between an original study and a replication study is higher than the permuted pattern similarity.
- **Classification tests for pattern expression values** An *a priori* pattern-based model from an original study can be used to examine whether the *a priori* model can classify the target conditions in a replication study.

## Acknowledgements

This work was supported by IS-R015-D1 (Institute for Basic Science), 2019R1C1C1004512 (National Research Foundation of Korea) (C.-W.W) and NIH R01DA035484, R01MH076136 (T.D.W.). The authors have no conflicts of interest to declare.

## Reference

Alink, A., Krugliak, A., Walther, A., Kriegeskorte, N., 2013. fMRI orientation decoding in V1 does not require global maps or globally coherent orientation stimuli. Frontiers in psychology 4, 493.

Ashar, Y.K., Chang, L.J., Wager, T.D., 2017. Brain Mechanisms of the Placebo Effect: An Affective Appraisal Account. Annu Rev Clin Psychol 13, 73–98.

Baker, M., 2016. 1,500 scientists lift the lid on reproducibility. Nature News 533, 452.

Behrens, T.E., Fox, P., Laird, A., Smith, S.M., 2013. What is the most interesting part of the brain? Trends Cogn Sci 17, 2–4.

Botvinick, M., Nystrom, L.E., Fissell, K., Carter, C.S., Cohen, J.D., 1999. Conflict monitoring versus selection-for-action in anterior cingulate cortex. Nature 402, 179–181.

Brabec, J., Rulseh, A., Hoyt, B., Vizek, M., Horinek, D., Hort, J., Petrovicky, P., 2010. Volumetry of the human amygdala-an anatomical study. Psychiatry Res 182, 67–72.

Brett, M., Christoff, K., Cusack, R., Lancaster, J., 2001. Using the Talairach atlas with the MNI template. Neuroimage 13, S85.

Bürkner, P.-C., 2017. brms: An R Package for Bayesian Multilevel Models Using Stan. Journal of Statistical Software 80, 1–28.

Button, K.S., Ioannidis, J.P., Mokrysz, C., Nosek, B.A., Flint, J., Robinson, E.S., Munafo, M.R., 2013. Power failure: why small sample size undermines the reliability of neuroscience. Nat Rev Neurosci 14, 365–376.

Carp, J., 2012. On the plurality of (methodological) worlds: estimating the analytic flexibility of FMRI experiments. Front Neurosci 6, 149.

Carpenter, B., Gelman, A., Hoffman, M.D., Lee, D., Goodrich, B., Betancourt, M., Brubaker, M., Guo, J., Li, P., Riddell, A., 2017. Stan: A probabilistic programming language. Journal of Statistical Software 76.

Collaboration, O.S., 2015. Estimating the reproducibility of psychological science. Science 349, aac4716.

Cremers, H.R., Wager, T.D., Yarkoni, T., 2017. The relation between statistical power and inference in fMRI. PLoS One 12, e0184923.

Eisenberger, N.I., Lieberman, M.D., Williams, K.D., 2003. Does rejection hurt? An fMRI study of social exclusion. Science 302, 290–292.

Gabrieli, J.D., Ghosh, S.S., Whitfield-Gabrieli, S., 2015. Prediction as a humanitarian and pragmatic contribution from human cognitive neuroscience. Neuron 85, 11–26.

Gelman, A., Hill, J., 2007. Data Analysis Using Regression and Multilevel/Hierarchical Models. Cambridge University Press.

Gilron, R., Rosenblatt, J., Koyejo, O., Poldrack, R.A., Mukamel, R., 2017. What’s in a pattern? Examining the type of signal multivariate analysis uncovers at the group level. Neuroimage 146, 113–120.

Gorgolewski, K.J., Varoquaux, G., Rivera, G., Schwarz, Y., Ghosh, S.S., Maumet, C., Sochat, V.V., Nichols, T.E., Poldrack, R.A., Poline, J.-B., 2015. NeuroVault. org: a web-based repository for collecting and sharing unthresholded statistical maps of the human brain. Frontiers in neuroinformatics 9, 8.

Haxby, J.V., Connolly, A.C., Guntupalli, J.S., 2014. Decoding neural representational spaces using multivariate pattern analysis. Annual review of neuroscience 37, 435–456.

Haxby, J.V., Guntupalli, J.S., Connolly, A.C., Halchenko, Y.O., Conroy, B.R., Gobbini, M.I., Hanke, M., Ramadge, P.J., 2011. A common, high-dimensional model of the representational space in human ventral temporal cortex. Neuron 72, 404–416.

Hayden, B.Y., Heilbronner, S.R., Pearson, J.M., Platt, M.L., 2011. Surprise signals in anterior cingulate cortex: neuronal encoding of unsigned reward prediction errors driving adjustment in behavior. J Neurosci 31, 4178–4187.

Hutson, M., 2018. Artificial intelligence faces reproducibility crisis. American Association for the Advancement of Science.

Ioannidis, J.P., 2005. Why most published research findings are false. PLoS Med 2, e124.

Ito, S., Stuphorn, V., Brown, J.W., Schall, J.D., 2003. Performance monitoring by the anterior cingulate cortex during saccade countermanding. Science 302, 120–122.

Johnson, R.A., Wichern, D.W., 2007. Applied Multivariate Statistical Analysis. Pearson Prentice Hall.

Kamitani, Y., Tong, F., 2005. Decoding the visual and subjective contents of the human brain. Nature neuroscience 8, 679.

Kang, J., Johnson, T.D., Nichols, T.E., Wager, T.D., 2011a. Meta Analysis of Functional Neuroimaging Data via Bayesian Spatial Point Processes. J Am Stat Assoc 106, 124–134.

Kang, J., Johnson, T.D., Nichols, T.E., Wager, T.D., 2011b. Meta analysis of functional neuroimaging data via Bayesian spatial point processes. Journal of the American Statistical Association 106, 124–134.

Kober, H., Barrett, L.F., Joseph, J., Bliss-Moreau, E., Lindquist, K., Wager, T.D., 2008. Functional grouping and cortical-subcortical interactions in emotion: a meta-analysis of neuroimaging studies. Neuroimage 42, 998–1031.

Kolling, N., Wittmann, M.K., Behrens, T.E., Boorman, E.D., Mars, R.B., Rushworth, M.F., 2016. Value, search, persistence and model updating in anterior cingulate cortex. Nat Neurosci 19, 1280–1285.

Kragel, P.A., Kano, M., Van Oudenhove, L., Ly, H.G., Dupont, P., Rubio, A., Delon-Martin, C., Bonaz, B.L., Manuck, S.B., Gianaros, P.J., 2018a. Generalizable representations of pain, cognitive control, and negative emotion in medial frontal cortex. Nature neuroscience, 1.

Kragel, P.A., Koban, L., Barrett, L.F., Wager, T.D., 2018b. Representation, Pattern Information, and Brain Signatures: From Neurons to Neuroimaging. Neuron 99, 257–273.

Kriegeskorte, N., 2009. Relating population-code representations between man, monkey, and computational models. Frontiers in Neuroscience 3, 35.

Kriegeskorte, N., Kievit, R.A., 2013. Representational geometry: integrating cognition, computation, and the brain. Trends Cogn Sci 17, 401–412.

Krishnan, A., Woo, C.-W., Chang, L.J., Ruzic, L., Gu, X., López-Solà, M., Jackson, P.L., Pujol, J., Fan, J., Wager, T.D., 2016. Somatic and vicarious pain are represented by dissociable multivariate brain patterns. Elife 5.

Kvitsiani, D., Ranade, S., Hangya, B., Taniguchi, H., Huang, J., Kepecs, A., 2013. Distinct behavioural and network correlates of two interneuron types in prefrontal cortex. Nature 498, 363.

Lindquist, K.A., Wager, T.D., Kober, H., Bliss-Moreau, E., Barrett, L.F., 2012. The brain basis of emotion: a meta-analytic review. Behav Brain Sci 35, 121–143.

Munafò, M.R., Nosek, B.A., Bishop, D.V.M., Button, K.S., Chambers, C.D., Sert, N.P.d., Simonsohn, U., Wagenmakers, E.-J., Ware, J.J., Ioannidis, J.P.A., 2017. A manifesto for reproducible science. Nature Human Behaviour 1, 0021.

Nord, C.L., Valton, V., Wood, J., Roiser, J.P., 2017. Power-up: a reanalysis of ‘power failure’in neuroscience using mixture modelling. Journal of Neuroscience, 3592–3516.

Ochsner, K.N., Gross, J.J., 2005. The cognitive control of emotion. Trends Cogn Sci 9, 242–249.

Park, S.H., Russ, B.E., McMahon, D.B., Koyano, K.W., Berman, R.A., Leopold, D.A., 2017. Functional subpopulations of neurons in a macaque face patch revealed by single-unit fMRI mapping. Neuron 95, 971–981.e975.

Peelen, M.V., Wiggett, A.J., Downing, P.E., 2006. Patterns of fMRI activity dissociate overlapping functional brain areas that respond to biological motion. Neuron 49, 815–822.

Pernet, C., Poline, J.-B., 2015. Improving functional magnetic resonance imaging reproducibility. Gigascience 4, 15.

Reddan, M.C., Lindquist, M.A., Wager, T.D., 2017. Effect Size Estimation in Neuroimaging. JAMA Psychiatry 74, 207–208.

Rouder, J.N., Morey, R.D., Speckman, P.L., Province, J.M., 2012. Default Bayes factors for ANOVA designs. Journal of Mathematical Psychology 56, 356–374.

Rouder, J.N., Speckman, P.L., Sun, D., Morey, R.D., Iverson, G., 2009. Bayesian t tests for accepting and rejecting the null hypothesis. Psychonomic bulletin & review 16, 225–237.

Samartsidis, P., Montagna, S., Nichols, T.E., Johnson, T.D., 2017. The coordinate-based metaanalysis of neuroimaging data. Statistical science: a review journal of the Institute of Mathematical Statistics 32, 580.

Shackman, A.J., Salomons, T.V., Slagter, H.A., Fox, A.S., Winter, J.J., Davidson, R.J., 2011. The integration of negative affect, pain and cognitive control in the cingulate cortex. Nat Rev Neurosci 12, 154–167.

Shmuel, A., Chaimow, D., Raddatz, G., Ugurbil, K., Yacoub, E., 2010. Mechanisms underlying decoding at 7 T: ocular dominance columns, broad structures, and macroscopic blood vessels in V1 convey information on the stimulated eye. Neuroimage 49, 1957–1964.

Simmons, J.P., Nelson, L.D., Simonsohn, U., 2011. False-positive psychology: undisclosed flexibility in data collection and analysis allows presenting anything as significant. Psychol Sci 22, 1359–1366.

Swisher, J.D., Gatenby, J.C., Gore, J.C., Wolfe, B.A., Moon, C.-H., Kim, S.-G., Tong, F., 2010. Multiscale pattern analysis of orientation-selective activity in the primary visual cortex. Journal of Neuroscience 30, 325–330.

Szucs, D., Ioannidis, J.P., 2017. Empirical assessment of published effect sizes and power in the recent cognitive neuroscience and psychology literature. PLoS Biol 15, e2000797.

Turner, B.O., Paul, E.J., Miller, M.B., Barbey, A.K., 2018. Small sample sizes reduce the replicability of task-based fMRI studies. Communications Biology 1, 62.

Wager, T.D., Atlas, L.Y., Botvinick, M.M., Chang, L.J., Coghill, R.C., Davis, K.D., Iannetti, G.D., Poldrack, R.A., Shackman, A.J., Yarkoni, T., 2016. Pain in the ACC? Proc Natl Acad Sci U S A 113, E2474–2475.

Wager, T.D., Atlas, L.Y., Lindquist, M.A., Roy, M., Woo, C.-W., Kross, E., 2013. An fMRI-based neurologic signature of physical pain. New England Journal of Medicine 368, 1388–1397.

Wager, T.D., Kang, J., Johnson, T.D., Nichols, T.E., Satpute, A.B., Barrett, L.F., 2015. A Bayesian model of category-specific emotional brain responses. PLoS computational biology 11, e1004066.

Wager, T.D., Lindquist, M.A., Nichols, T.E., Kober, H., Van Snellenberg, J.X., 2009. Evaluating the consistency and specificity of neuroimaging data using meta-analysis. Neuroimage 45, S210–221.

Woo, C.-W., Chang, L.J., Lindquist, M.A., Wager, T.D., 2017a. Building better biomarkers: brain models in translational neuroimaging. Nature neuroscience 20, 365.

Woo, C.-W., Schmidt, L., Krishnan, A., Jepma, M., Roy, M., Lindquist, M.A., Atlas, L.Y., Wager, T.D., 2017b. Quantifying cerebral contributions to pain beyond nociception. Nature communications 8, 14211.

Woo, C.W., Koban, L., Kross, E., Lindquist, M.A., Banich, M.T., Ruzic, L., Andrews-Hanna, J.R., Wager, T.D., 2014a. Separate neural representations for physical pain and social rejection. Nat Commun 5, 5380.

Woo, C.W., Krishnan, A., Wager, T.D., 2014b. Cluster-extent based thresholding in fMRI analyses: pitfalls and recommendations. Neuroimage 91, 412–419.

Yarkoni, T., Poldrack, R.A., Nichols, T.E., Van Essen, D.C., Wager, T.D., 2011. Large-scale automated synthesis of human functional neuroimaging data. Nat Methods 8, 665–670.

